# Correlative three-dimensional fluorescence and refractive index tomography: bridging the gap between molecular specificity and quantitative bioimaging

**DOI:** 10.1101/186734

**Authors:** Kyoohyun Kim, Wei Sun Park, Sangchan Na, Sangbum Kim, Taehong Kim, Won Do Heo, YongKeun Park

## Abstract

Optical diffraction tomography (ODT) provides label-free three-dimensional (3D) refractive index (RI) measurement of biological samples. However, due to the nature of the RI values of biological specimens, ODT has limited access to molecular specific information. Here, we present an optical setup combining ODT with three-channel 3D fluorescence microscopy, to enhance the molecular specificity of the 3D RI measurement. The 3D RI distribution and 3D deconvoluted fluorescence images of HeLa cells and NIH-3T3 cells are measured, and the cross-correlative analysis between RI and fluorescence of live cells are presented.

## 1. Introduction

The three-dimensional (3D) imaging of biological cells and tissues has been an invaluable tool for investigating subcellular organelles and various biological mechanisms [1]. Among them, optical diffraction tomography (ODT) or holotomography (HT) has emerged as an effective live-cell imaging technique [2, 3]. ODT reconstructs the 3D refractive index (RI) distribution of a sample. Because RI is an intrinsic optical property of materials, ODT enables the 3D imaging of live cells and tissues without using any exogenous labeling agents or preparation steps [4-9]. This is important because the use of fluorescence proteins, quantum dots, or dye molecules inevitably introduces unwanted effects such as phototoxicity or photobleaching, and in some cases cells should be fixed for measurements [10]. Furthermore, since the RI value of most biological specimen is linearly proportional to protein concentration [11, 12], ODT provides quantitative information including protein concentration and dry mass. For these reasons, there are growing numbers of biological and medical studies employing ODT to investigate the physiology of various samples [3] including neuron cells [13, 14], red blood cells [15-17], parasites in host cells [18, 19], immune cells [20], bacteria [13, 21], gold nanoparticles in live cells [22], embryos [23], live cells in a microfluidic channel [24-26], and blood cells *in vivo* [27].

While the measured 3D RI distribution provides the label-free analysis of cells and subcellular organelles, ODT offers limited molecular specificity. For instance, ODT can measure the protein concentration and dry mass of eukaryotic cells containing a variety of proteins, because most proteins have similar RI increment values [28]. However, since the RI value itself does not provide molecular specific information which can be used to identify proteins and subcellular organelles, it is difficult to distinguish a specific protein inside each intracellular component based on the measured RI distribution.

Recently, several approaches have been proposed to enhance the molecular specificity of ODT. For example, the RI dispersion of samples was utilized [29]. In addition, the segmentation with RI values has provided molecular specific information of distinct materials such as lipid droplets [30], polystyrene beads [20], and gold nanoparticles inside cells [22]. More recently, multimodal approaches have been demonstrated. Combining structured illumination microscopy (SIM) [31, 32] with ODT was demonstrated for addressing sub-diffraction-resolution molecular imaging [33]. In addition, an integrated setup for fluorescence microscopy and ODT with a sample rotation scheme showed enhanced molecular specificity and an isotropic spatial resolution for 3D RI tomography [34].

In this paper, we present a method combining ODT with three-channel 3D fluorescence microscopy. Exploiting a Mach-Zehnder interferometry equipped with a dynamic micromirror device (DMD), 3D RI maps of a sample is measured with high speed and precision. Epi-fluorescence microscopy system is integrated into this system in order to simultaneously measured multi-channel 3D fluorescence image of the sample. To demonstrate the capability of the method, both the 3D RI tomograms and 3D fluorescence images were measured for HeLa cells and NIH-3T3 cells. Our results show that the RI values of cell nuclei could be either higher or lower than surrounding cytoplasm, depending on cell types.

## 2. Methods

### 2.1 Optical setup

The setup consists of ODT and epi-fluorescence microscopy in the same optical imaging system (Fig. 1). In order to measure the 3D RI distribution of a cell, ODT employing Mach-Zehnder interferometry was used [Fig. 1(a)]. A 532-nm diode-pumped solid-state laser beam (MSL-S-532-10mW, CNI laser, China) was coupled into a 2×2 fiber coupler (FUSED-12-532-3.5, OZoptics, Canada), and was split into two arms: a sample and a reference arm. The sample illumination beam was reflected by a DMD (DLP6500FYE, Texas Instruments, USA. The DMD diffracted the beam into several orders, among which the first-order diffracted beam was collected by a tube lens and a condenser lens [Numerical aperture (NA) = 0.7, 60×] to impinge onto the sample. The diffracted beam scattered by the sample was collected by an objective lens (NA = 0.8, 60×) and a tube lens, and interfered with the reference beam to generate a spatially modulated hologram at a camera plane.

**Figure 1.**
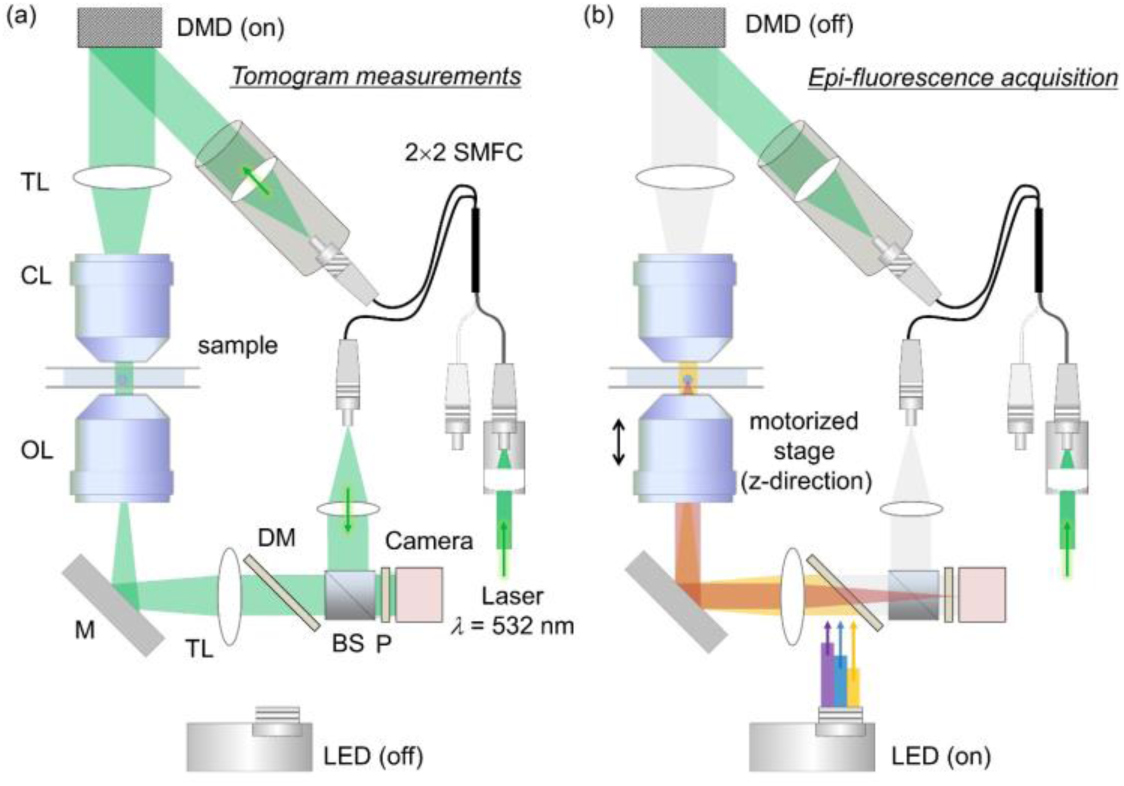
The combined optical setup for (a) optical diffraction tomography and (b) 3D epi-fluorescence microscopy. SMFC: single-mode fiber coupler, DMD: digital micromirror device, TL: tube lens, CL: condenser lens, OL: objective lens, M: mirror, DM: dichroic mirror, BS: beam splitter, P: polarizer, LED: laser emitting diode.

For tomographic reconstruction, the illumination beam was circularly scanned with plane waves with 49 different azimuthal angles. To control the incident angle of the illumination beam, the DMD displays a 4-bit-depth amplitude grating pattern, in which a spatial groove corresponds to the spatial frequency of the tilted illumination beam. Each gray-scale amplitude grating pattern was generated by time-multiplexing four binary patterns, e.g., the Lee holograms [35, 36]. By displaying the four binary patterns sequentially during the camera exposure, unwanted diffraction patterns from the binary patterns were eliminated by time-multiplexed averaging [37]. A high-speed DMD and camera were employed, and tomogram acquisition was executed within 0.4 sec.

3D fluorescence imaging was performed using the same optical setup [Fig. 1(b)]. Epifluorescence geometry was used for the excitation of the fluorescence probes and the measurements of emission signals. In order to excite fluorescence probes inside a sample, a multi-wavelength light-emitting diode (LED) source (center wavelength: *λ*_B_ = 385 nm, *λ*_G_ = 470 nm, *λ*_R_ = 565 nm) was coupled to the optical setup using a three-channel dichroic mirror (Semrock FF409/493/596-Di01). During the fluorescence image acquisition, the DMD was turned off to avoid camera saturation and potential photobleaching by the laser beam.

### 2.2 Optical diffraction tomography algorithm

Figures 2(a)–(e) show the procedure for 3D RI tomogram reconstruction using ODT. The ODT algorithm is an optical analogy to X-ray computed tomography. The 2D optical fields of a sample (NIH-3T3 cell) are retrieved from spatially modulated interferograms *I*(*x*, *y*) obtained with various illumination angles [Fig. 2(a)]. From the recorded interferograms, complex 2-D optical fields, consisting of both the amplitude |*U*| and the phase images ∠*U* of the sample were retrieved by a field retrieval algorithm [38, 39]. Then from the multiple 2D optical fields, ODT reconstructs a 3D RI tomogram of the sample *n*(*x*, *y*, *z*) by inversely solving light diffraction at a weakly scattering sample [2], as shown in Fig. 2(d). For the purpose of visualization, pseudo coloring was applied based on the RI values [40]. Because of the limited NAs for the condenser and objective lenses, a fraction of the light diffracted from the sample is not collected, resulting in missing information in the reconstruction. To address this issue, known as the missing cone problem, an iterative approach with the non-negativity constraint was used as a regularization algorithm [41]. The detailed information for the ODT system and the tomogram reconstruction algorithms can be found elsewhere [40, 42].

**Figure 2.**
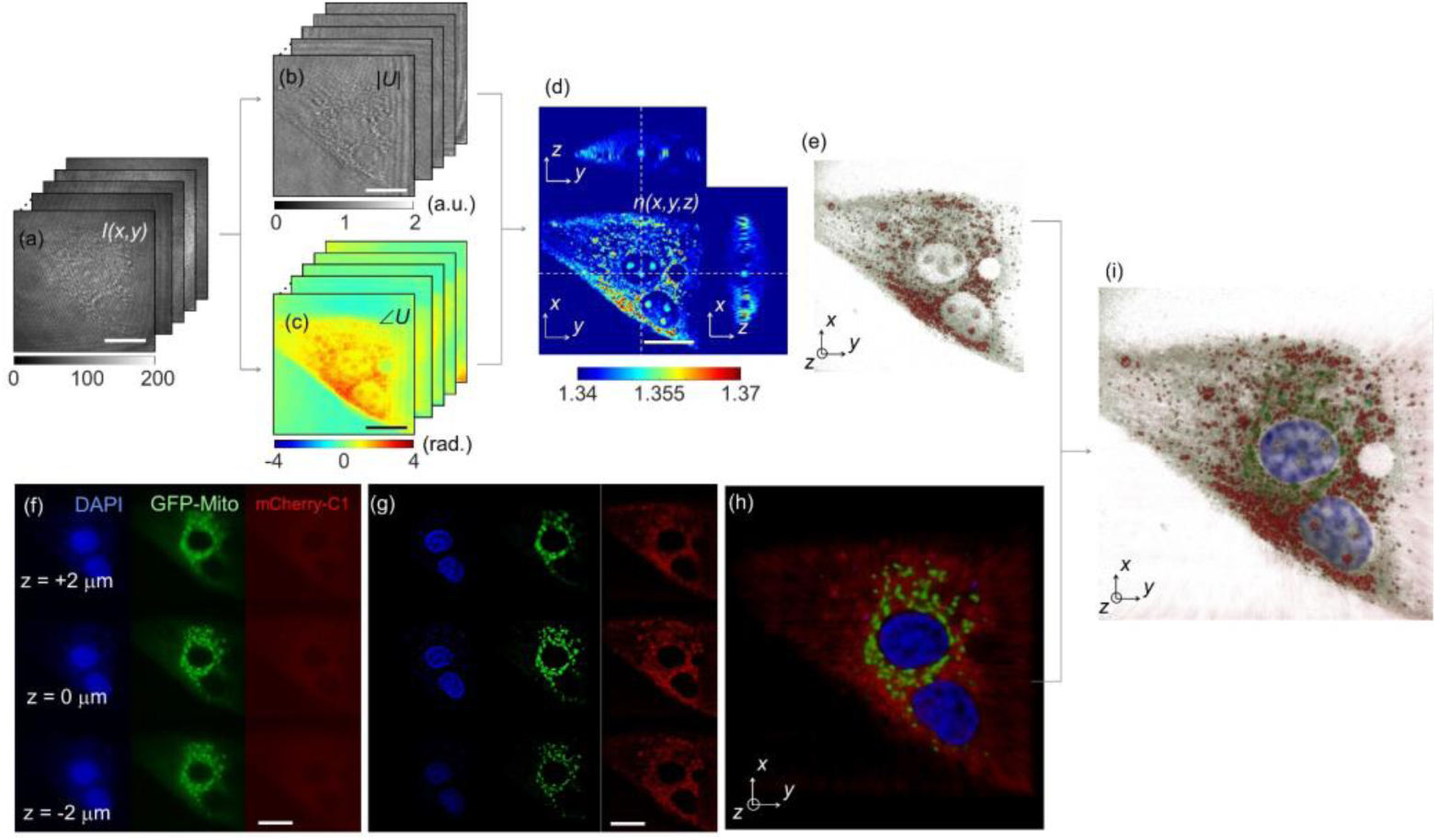
A schematic diagram of the ODT reconstruction (a-e) and correlative microscopy with 3D fluorescence imaging (f-h). (a) Raw holograms of an NIH-3T3 cell obtained with various illumination angles. (b) Amplitude and (c) phase map of the complex optical fields of the cell. (d) The cross-sectional slices and (e) the 3D rendered map of the reconstructed 3D RI distribution of the sample. (f) Axially stacked three-channel epi-fluorescence images of the same cell. (g) The fluorescence images of the cell after 3D deconvolution. (h) The fluorescence image merged with three-channel deconvoluted fluorescence images. (i) The 3D rendered map of the 3D RI distribution and three-channel fluorescence images for correlative analysis. Scale bar indicates 20 μm.

### 2.3 3D fluorescence image acquisition

To acquire 3D fluorescence images, the sample stage was scanned along the axial direction with a step size of 469 nm and a scanning distance of 15 μm. At each axial position, the LED source switched the excitation wavelength of the LED, and measured multi-channel emission signals sequentially. Typically, the camera was exposed for 1 s to acquire a single-channel fluorescence image at one axial position.

Figures 2(f)–(h) show the procedures for the 3D multichannel fluorescence microscopy. The same NIH-3T3 cell used in Figs. 2(a)–(e) was imaged using the 3D fluorescence imaging mode. The nucleus, mitochondria, and actin are labeled with DAPI, green fluorescence proteins (GFP-Mito), and mCherry-C1, respectively, and excited with three LED sources with different spectra (center wavelengths are 385 nm, 470 nm, and 565 nm), respectively. Figure 2(f) shows the acquired 2D fluorescence images of the sample at the focal position of *z* = 2 μm, 0 μm, and −2 μm.

In order to enhance the image contrast and the spatial resolution of the fluorescence images, 3D deconvolution was performed using commercial software (Huygens Professional, Scientific Volume Imaging b.v., the Netherlands), which exploits the iterative classic maximum likelihood estimation. The measured axial stacks of each fluorescence channel were deconvoluted using theoretical 3D point spread functions of the optical setup, which were calculated from optical parameters including the wavelength of the emission signals and the numerical aperture of the objective lens. As shown in Fig. 2(g), the deconvoluted fluorescence images show high image contrast compared to the raw epi-fluorescence images in Fig. 2(f). Then, the three-channel deconvoluted fluorescence images were merged in three-dimensions. The obtained fluorescence images of the sample in three channels were then merged into a single combined fluorescence image [Fig. 2(h)], which can also be correlated with the independently measured 3D RI tomogram of the same sample [Fig. 2(i)].

### 2.4 Sample preparation

HeLa (ATCC CCL 2) cells and NIH-3T3 (ATCC CRL 1658) cells were purchased from ATCC and maintained in Dulbecco’s Modified Eagle’s Medium (DMEM; High Glucose, Pyruvate, Gibco), supplemented with 10% FBS (Invitrogen) and 1% Penicillin-Streptomycin (10,000 U/mL) at 37°C in a humidified 10% CO_2_ atmosphere.

HeLa cells and NIH-3T3 cells were transfected with DNA plasmids (mCherry-C1 and GFP-Tubulin) using Lipofectamine LTX DNA Transfection Reagent (#15338-100) in a 12-well dish first and then the cells were subcultured in a culture dish (Tomodish, Tomocube Inc., Republic of Korea). The cells were stained with Hoechst (0.1 (μg/ml) and washed with fresh growth medium prior to imaging.

## 3. Results

### 3.1 Combined 3D RI and 3D fluorescence images

Figure 3 presents the combined 3D RI and deconvoluted fluorescence images of HeLa cells and an NIH-3T3 cell. The 3D RI distribution of cells clearly shows both the overall cellular morphology and the subcellular structures including the nucleus, nucleoli, and intracellular vesicles. The three-channel deconvoluted fluorescence images provide high molecular specific images, presenting nuclei (blue), tubulin (green), and actin (red) inside the cells separately. In Fig. 3, individual cytoskeleton structures are not distinguishable, which might have resulted from the movements of live cells.

Since ODT and 3D deconvolution fluorescence microscopy employ the same optics setup, the 3D RI distribution and fluorescence images of cells are well-matched, which enables the identification of each subcellular component from the label-free 3D RI distribution of the samples. For instance, the blue-channel fluorescence images clearly exhibit interior nucleus cells, which can also be identified by a visual inspection of the 3D RI distribution of cells.

### 3.2 Cross-correlative analysis of 3D RI and 3D fluorescence images

The present technique provides the cross-correlative analysis of 3D RI and 3D deconvoluted fluorescence images which enables localization of the 3D RI values of each subcellular region. The reconstructed 3D RI distributions of HeLa cells and an NIH-3T3 cell in Figs. 4(a)–(b) and Figs. 4(d)–(e), respectively, visually outline their internal structures including nucleus regions, nucleoli, and cytoplasmic regions inside the cells, while it can only be confirmed by comparing with the 3D deconvoluted fluorescence images in Fig. 4(c) and 4(f). Here, the nucleus regions stained with Hoechst can be identified in the blue channel, while cytoplasmic regions where most tubulin networks are located can be found in the green channel.

**Figure 3.**
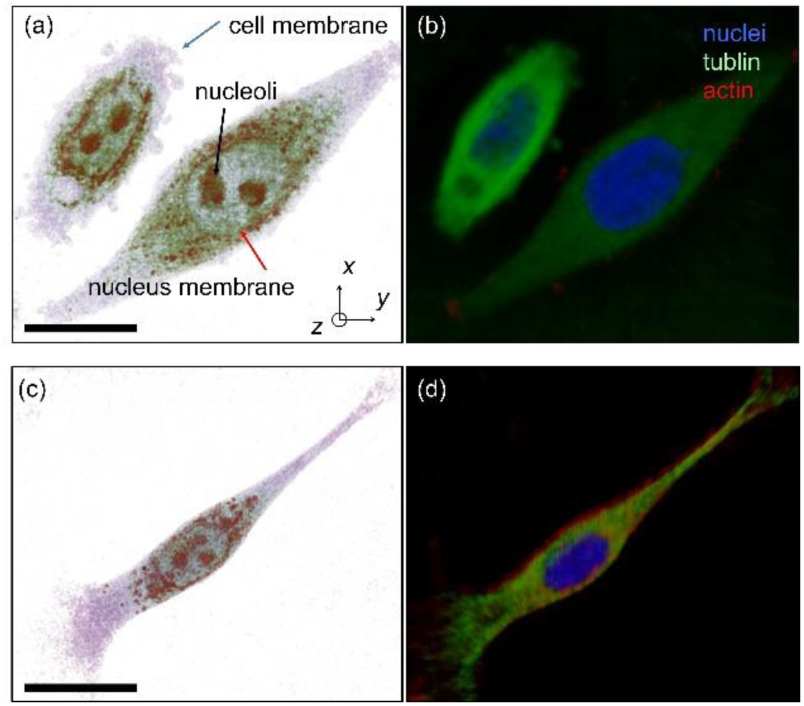
(a, c) The 3D rendered map of the 3D RI distribution and (b, d) 3D deconvoluted three-channel fluorescence image of a HeLa cell and an NIH-3T3 cell, respectively. Scale bar indicates 20 μm.

**Figure 4.**
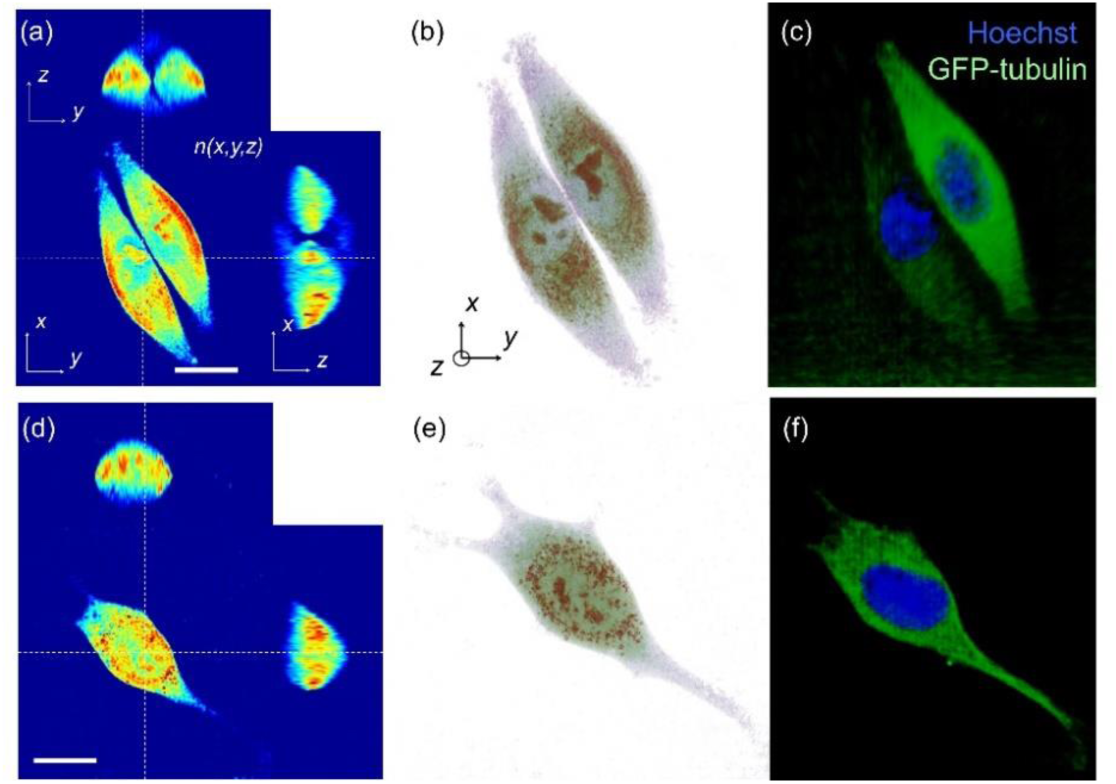
Cross-correlative analysis of 3D RI and 3D fluorescence images of (a–c) a HeLa cell and (d–f) an NIH-3T3 cell. (a, e) The cross-sectional slices and (b, f) the 3D rendered map of reconstructed 3D RI distribution of the samples. Scale bar indicates 20 μm. (c, g) The 3D deconvoluted fluorescence images of the samples stained with Hoechst (blue) and GFP-tubulin (green).

From the measured 3D RI and 3D deconvoluted fluorescence images, the 3D RI distributions of each nucleus and cytoplasmic region were measured separately. In the 3D deconvoluted fluorescence images, the nucleus and cytoplasmic regions of each cell were segmented by voxels which had fluorescence signals higher than the background values in the blue and green channel fluorescence images, respectively. Then, the segmented subcellular regions in the 3D fluorescence images were employed as filtering masks in the RI tomograms of the cells, which provide the 3D RI distribution of each subcellular region.

To demonstrate the potential of the correlative images, we compared the RI values of the nucleus of the HeLa and the NIH-3T3 cell. In the HeLa cell, the overall values of the nucleus were smaller than that of the cytoplasm. Although the RI values of the nucleolus are compatible to that of the cytoplasm, the RI values of the nucleoplasm including chromatin are significantly smaller than the average RI value of the cytoplasm. However, the NIH-3T3 cell shows that the RI values of nucleoplasm are compatible to that of the cytoplasm, while the RI value of the nucleoli is higher than the average value of the nucleoplasm.

### 3.3 RI value of the nucleoplasm

To further analyze the RI value of the nucleus, the RI histograms of the nucleus and cytoplasmic regions in HeLa cells (*n* = 4) and NIH-3T3 cells (*n* = 6) are shown in Fig. 5(a) and 5(b), respectively. As shown in the cases of individual cell images (Fig. 4), the RI values of the nucleoplasm and nucleus of most of the HeLa cells are lower the average values of cytoplasm, whereas those of the NIH 3T3 cells are higher than that of cytoplasm.

**Figure 5.**
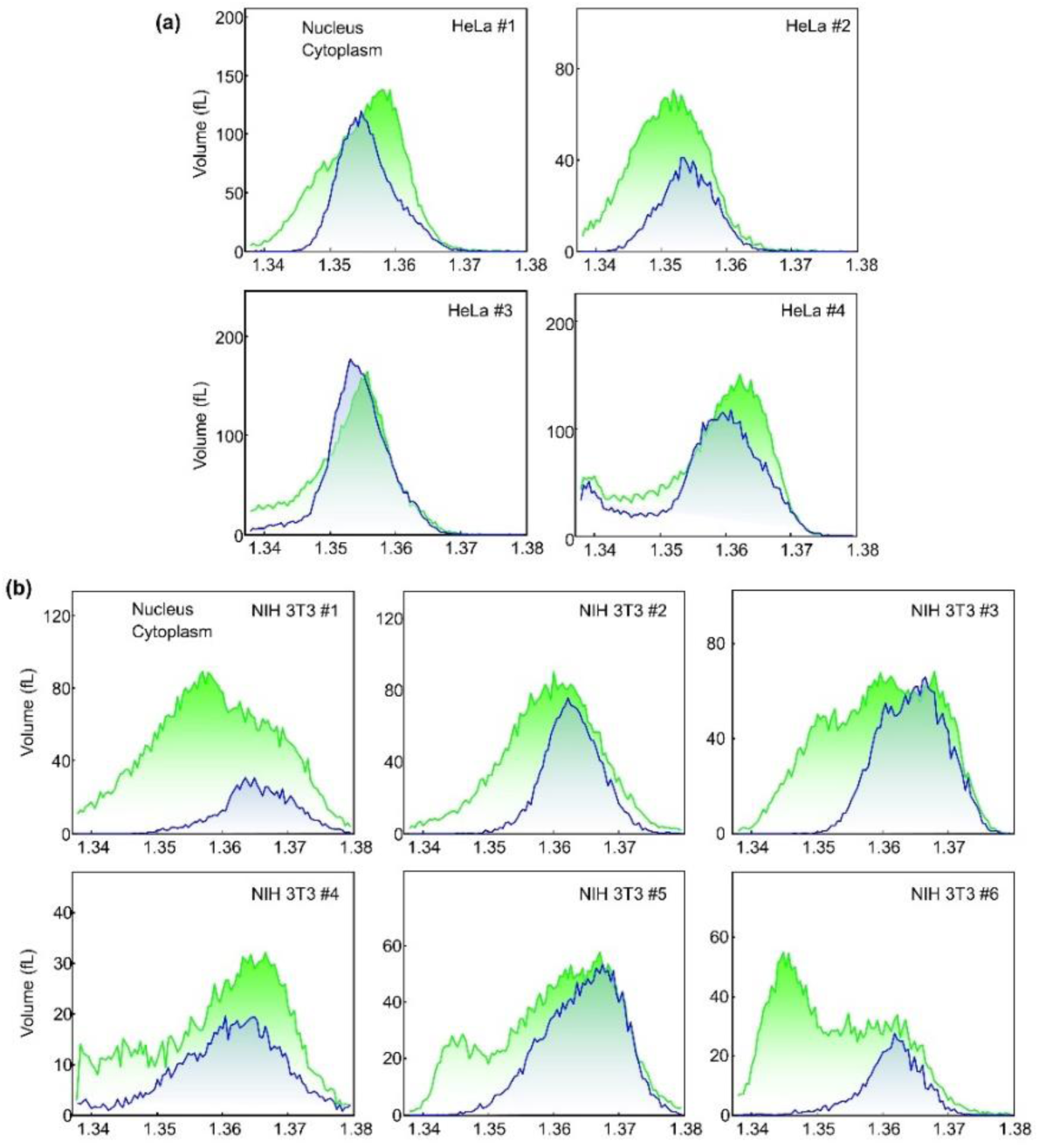
Histograms of RI values in the nucleus and cytoplasm of HeLa cells (a) and NIH-3T3 cells (b) (*n* = 4 for HeLa cells, *n* = 6 for NIH-3T3 cells)

Previously, a body of papers reported that the RI values of nuclei are lower than the surrounding cytoplasm [34, 43-45], whereas several other papers provide results indicating the RI values of the nuclei are higher than the cytoplasm [46-48]. This series of papers in two groups seem to contact with each other. However, our results show that RI values of nuclei can be lower or higher than surrounding cytoplasm. The results clearly show that whether RI values of nuclei is higher or lower than the surrounding cytoplasm can be affected by the types of cell.

Furthermore, the high-resolution measurements of 3D RI and 3D fluorescence images also indicate that even in nuclei significant spatial variation in RI values occurs, as shown in Fig. 4. In particular, the chromatin structures exhibit higher RI values than the nucleoplasm in both the NIH-3T3 and Hela cells.

## 4. Discussion and Conclusion

Here, we present a method for measuring both the 3D RI distribution and three-channel 3D fluorescence image of live cells. Employing a combined optical setup for ODT and epi-fluorescence microscopy, 3D RI tomograms and three-channel 3D deconvoluted fluorescence images of biological cells were measured with high spatial resolution. Since both microscopic imaging modalities share the same optical setup, the present approach enables direct measurement and precise analysis for correlative imaging. Like PET/CT provides both the morphological and molecule specific imaging in medical diagnosis, the present method is an optical analogy which provides both morphological and molecular imaging of biological cells. To demonstrate the capability of the method, both 3D RI tomograms and 3D fluorescence images of HeLa cells and NIH-3T3 cells were obtained. Our results clearly demonstrate that the use of both 3D RI and fluorescence images provide comprehend and also complementary information about cell physiology. In addition, our results show that the RI values of cell nuclei can be either higher or lower than the surrounding cytoplasm, depending on the cell types.

As the experimental results show, the addition of 3D fluorescence images provides molecular specific information about the live cells, together with the morphological, biophysical, and biomechanical properties that ODT provides. Furthermore, the ability to measure three-channel 3D fluorescence information can provide useful information when correlated with RI tomograms of the cell. These two different modalities are complementary to each other, and will allow various biological studies where the systematic investigation of subcellular structures and their dynamics are critical.

This new instrument has several potential applications, one of which we explain here; tracking and imaging subcellular organelles such as mitochondria or chromatins for a long period of time. An initial measurement using 3D fluorescence imaging enables the localization of specific subcellular structures exploiting molecular specificity. Then, consecutive measurements using dynamic 3D RI tomograms of live cells can allow tracking and imaging these target organelles over an extended time period with minimal exposure to the issues of phototoxicity and photodamaging. Furthermore, the ability to obtain fast time-lapse measurements (up to 100 tomograms per second) using ODT makes it possible to visualize biologically important dynamics which cannot be accessed by fluorescence imaging techniques [49].

The present instrument can be further improved to enhance acquisition speed. Currently, it takes approximately 1 second to acquire a single-channel fluorescence image at one axial position. Consequently, the total acquisition time for three-channel fluorescence images axially scanned with a step size of 469 nm and a scanning distance of 15 μm would take ~100 seconds, which might be an issue for fast dynamics measurements. The use of a sensitive and fast image sensor such as a scientific CMOS sensor would further enhance acquisition speed. In addition, the present method can also be combined with optical manipulation. By doing so, specific parts of live cells can be identified first with ODT and 3D fluorescence images, which are then optically controlled for other purposes [50, 51].

## Acknowledgement

This work was supported by KAIST, BK21+ program, Tomocube, and National Research Foundation of Korea (2015R1A3A2066550, 2014M3C1A3052567, 2014K1A3A1A09063027). Dr. K. Kim, Dr. W. S. Park, and Prof. Y. Park has financial interests in Tomocube Inc., a company that commercializes ODT and quantitative phase imaging instruments and is one of the sponsors of the work.

